# Chilean brush-tailed mouse (*Octodon degus*): a diurnal precocial rodent as a new model to study visual receptive field properties of superior colliculus neurons

**DOI:** 10.1101/2024.03.25.586655

**Authors:** N.I. Márquez, P.F. Fernández-Aburto, A. Deichler, I. Perales, J.-C. Letelier, G. J. Marín, J. Mpodozis, S.L. Pallas

## Abstract

Lab rodent species used to study the visual system and its development (hamsters, rats, and mice) are nocturnal, altricial, and possess simpler visual systems than carnivores and primates. To widen the spectra of studied species, here we introduce an alternative model, the Chilean degu (*Octodon degus*), a diurnal, precocial Caviomorph rodent with a cone enriched, well-structured retina, and well-developed central visual projections. To assess degus’ visual physiological properties, we characterized the visual responses and receptive field (RF) properties of isolated neurons in the superficial layers of the superior colliculus (sSC). To facilitate comparison with studies in other rodent species, we used four types of stimuli: (1) a moving white square, (2) sinusoidal gratings, (3) an expanding black circle (looming), and (4) a stationary black circle. We found that as in other mammalian species, RF size increases from superficial to deeper SC layers. Interestingly, compared to other lab rodents, degus have smaller RF sizes, likely indicating higher acuity. sSC neurons displayed spatial frequency tuning to grating stimuli from 0.08 to 0.24 cycles/degree. Additionally, neurons from sSC showed transient ON, OFF, or ON-OFF responses to stationary stimuli but increased their firing rates as a looming object increased in size. Our results suggests that degus have higher visual acuity, higher frequency tuning, and lower contrast sensitivity than commonly used nocturnal lab rodents, positioning degus as a well-suited model for studies of diurnal vision that are more relevant to humans.

## INTRODUCTION

In studies of visual processing circuitry, the choice of animal model is often based on tradition or convenience. In contrast, animals that have been chosen for auditory processing research are species that have exceptional auditory processing ability. It could easily be argued that more has been learned about auditory signal processing from studies of bats (Razak 2011; Razak and Fuzessery 2015), owls (DeBello and Knudsen 2004), songbirds (Brainard and Doupe 2013), and chinchillas (Jones *et al*. 2015), than from traditional lab animals. The foundational studies of central visual processing were done largely in cats and monkeys (Hubel and Wiesel 1998), but mice are now the model of choice. Although mouse visual system is similar to that of other mammals in many ways (Huberman and Niell 2011; Garrett *et al*. 2014; Juavinett and Callaway 2015), there are important differences, possibly related to their nocturnal habit and poor eyesight, that have unknown consequences for reaching a general understanding of visual system development and function.

In the Pallas lab, the superior colliculus (SC) and visual cortex (V1) of Syrian hamsters were chosen for studies of visual circuit development, largely because hamsters are born earlier and in larger litters than more typically chosen rodent subjects. As in other mammals studied, receptive fields (RFs) in hamster SC and V1 are large at birth and undergo a postnatal refinement process to reach adult size. It had been assumed, based largely on studies of primary visual cortex (V1) in cats and monkeys (Hensch *et al*. 1998; Espinosa and Stryker 2012), that visual pathway development requires early visual experience during a critical period for maturation but not for maintenance of refined receptive fields and response properties. In contrast, the Pallas lab has found that developmental refinement of receptive fields in SC and V1 occurs normally in dark reared hamsters. Continued visual deprivation in adulthood results in a gradual loss of RF refinement in both SC and V1 of adult hamsters (> postnatal day (P) 60) (Carrasco *et al*. 2005; Balmer and Pallas 2015). A brief, late juvenile exposure to light stabilizes receptive field size permanently despite continued visual deprivation, but visual experience after sexual maturity (P 60) has no effect (Carrasco and Pallas 2006; Balmer and Pallas 2015). These results raise the question of whether hamsters are unique or if other rodents, in particular those that are diurnal, would exhibit similar independence from visual stimulation during development.

Here we characterize the RF properties of SC neurons in Chilean degus (*Octodon degus*), a diurnal, precocial rodent species (Fulk 1976; Reynolds and Wright 1979; Long and Ebensperger 2010) with a cone enriched retina and a well-developed visual system (Vega-Zuniga *et al*. 2013). Unlike typical lab rodent species, degus open their eyes at P1 and exhibit visually guided behaviors shortly after birth, exposing their visual system to an early visual experience that could shape its maturation. Our results show that degus have clear diurnal characteristics and a better visual acuity in comparison to nocturnal lab rodents. Here we argue that degus are a promising model to study the role of visual experience in the maturation of the visual system and thus might be better suited to model the human visual system.

## METHODS

### Animals

A total of 10 animals from our colony were used in this study. Animals were housed in metal cages (50 x 40 x 35 cm) with wood shavings as previously described by Márquez *et al*. (2015) in a climate-controlled room under a 12:12 photoperiod in groups of 3-4 degus of the same sex. They were fed with rodent-chow Prolab® RMH 3000 (Purina LabDiet®, Inc.), Mazuri chinchilla diet (Mazuri® Exotic Animal Nutrition) and provided with water *ad libitum*. All animal procedures were approved by the *Comité Institucional de Cuidado y Uso Animal* (CICUA) of the University of Chile and the IACUC of the University of Massachusetts-Amherst and followed the National Institutes of Health (NIH) Guide for the Care and Use of Laboratory Animals. All procedures met additional standards of care established by SfN and IBRO.

### Histology and neuronal tracers

Some degus (n=2) were used only for neuroanatomical data collection. The labeling of retinal projections to the SC was performed under gaseous anesthesia using isoflurane (4% in medical oxygen at a flow rate of 100 cc/kg/min) complemented with diazepam (5 mg/kg) administrated IP. A glass pipette (tip diameter 15–20 μm) containing 1% cholera toxin subunit B (CTB, List Biological Laboratories, Campbell, CA) in 0.1 M phosphate buffer was injected intraocularly. After 5–7 days survival, the degus were sedated with gaseous isoflurane, placed in a deep state of anesthesia with ketamine/xylazine (ketamine 40-80 mg/kg, xylazine 5-10 mg/kg), and perfused intracardially by passing 200 ml of warm saline solution (0.9% in 0.1 M PB) followed by 200 ml of cold paraformaldehyde (PFA, 4% in 0.1 M PBS). The brains were extracted and postfixed overnight in 4% PFA at 4°C, and then transferred to 30% sucrose solution in 0.1 M PB until they sank, for cryoprotection. Brains were sectioned (60 μm), and a CTB immunoreaction assay was performed as described by Deichler *et al*. (2019).

For histological confirmation of electrode positioning during electrophysiological recording, degus were given an overdose of urethane (3 mg/kg, IP) and perfused intracardially as described above. The brains were postfixed overnight in 4% PFA at 4°C and then transferred to 30% sucrose for cryoprotection. Finally, brains were sectioned (60 μm), mounted, and Nissl-stained (cresyl violet, Merck, Darmstadt, Germany). Reconstruction of electrode penetrations in several experiments verified that our recordings were taken at intervals of 100 microns through a transect passing across layers of the SC, from the SGS into the SGI.

### *In vivo* recordings

#### Animal Preparation

Pre-op: Degus store wood shavings in their cheeks (Deichler, pers. comm.). Thus, to avoid aspiration of the shavings during anesthesia, they were taken from their cages and placed in a clean cage with paper towels the night before the experiment. Degus (n = 8) were fasted overnight after receiving Dexamethasone (1 mg/kg, SQ) to prevent cerebral edema. The morning of surgery, a second dose of Dexamethasone (1 mg/kg, SQ) was given along with atropine (0.1 mg/kg, SQ) to counteract bradycardia and apnea and to reduce mucosal secretions.

#### Surgery

Isoflurane (4% in medical oxygen at a flow rate of 100 cc/kg/min) was used for anesthesia induction, then diazepam (5 mg/kg) was administrated IP to induce sedation before degus were anesthetized using urethane (1.25 mg/kg, IP). If necessary, Doxapram was used (2 mg/kg SQ or sublingual) to stimulate respiration. Body temperature was kept between 36 °C and 37 °C throughout the experiment by using a thermo-regulated blanket (FHC, Inc.). To prevent asphyxia due to accumulation of bronchial secretions, degus were tracheostomized after the site was prepped by shaving the neck and chest, cleaning the area with ethanol and povidone, and inserting a tracheal tube. The scalp was shaved, and the head stabilized in a stereotaxic head holder. Then, the surgical site was cleaned with ethanol and povidone followed by a midline incision in the scalp. The muscles and periosteum overlying the skull were retracted, and a small craniotomy above the SC region was made with a surgical micro-drill. After unilaterally aspirating the cortex to expose the SC and covering the site with saline, electrophysiological recordings were obtained by using Epoxy-insulated tungsten microelectrodes (1–2 MOhms; FHC, Inc.) or a 16 Channel multi-electrode array (MEA) (∼1 MOhm; Microprobes for Life Science, Inc.) and a multichannel differential amplifier (Model 3600; AM Systems Inc.). A silver chloride wire was implanted in between the exposed bone and muscle as a reference electrode. Physiological data were sampled just before, during, and just after the stimulation at 10-20 kHz, with a band-pass of 3 Hz to 10 kHz, by using a standard PC with a 16-channel A/D converter board (National Instruments Corp). Data acquisition and offline analysis were performed using Igor Pro-8 (WaveMetrics, Inc.) or MATLAB (MathWorks, Inc) software.

### Visual stimulation

Visual stimuli were presented on an LCD monitor (Alienware 25 AW2518HF, 1920 x 1080 pixels, 60 Hz refresh rate) placed 32 cm from the degu’s eye. Stimulus parameters were controlled with a lab-designed program using the Python (Python Software Foundation) - based open-source application PsychoPy (v.3.1.3). A photodiode that detected the stimulus produced a 5-volt TTL signal that was recorded by the amplifier and used to synchronize stimuli and neural responses. Four types of visual stimuli were used:

#### Drifting squares

A white square (∼3° diameter) on a dark background was swept from the top to the bottom of the monitor over 4 s, with 1 s intervals between successive stimuli. The position of the square changed in an orderly manner along the horizontal axis of the screen at each sweep, covering the entire screen within 64 runs (**Fig. 1A**). The stimulation protocol was repeated 3 times for each recording location/depth.

**Figure 1.**
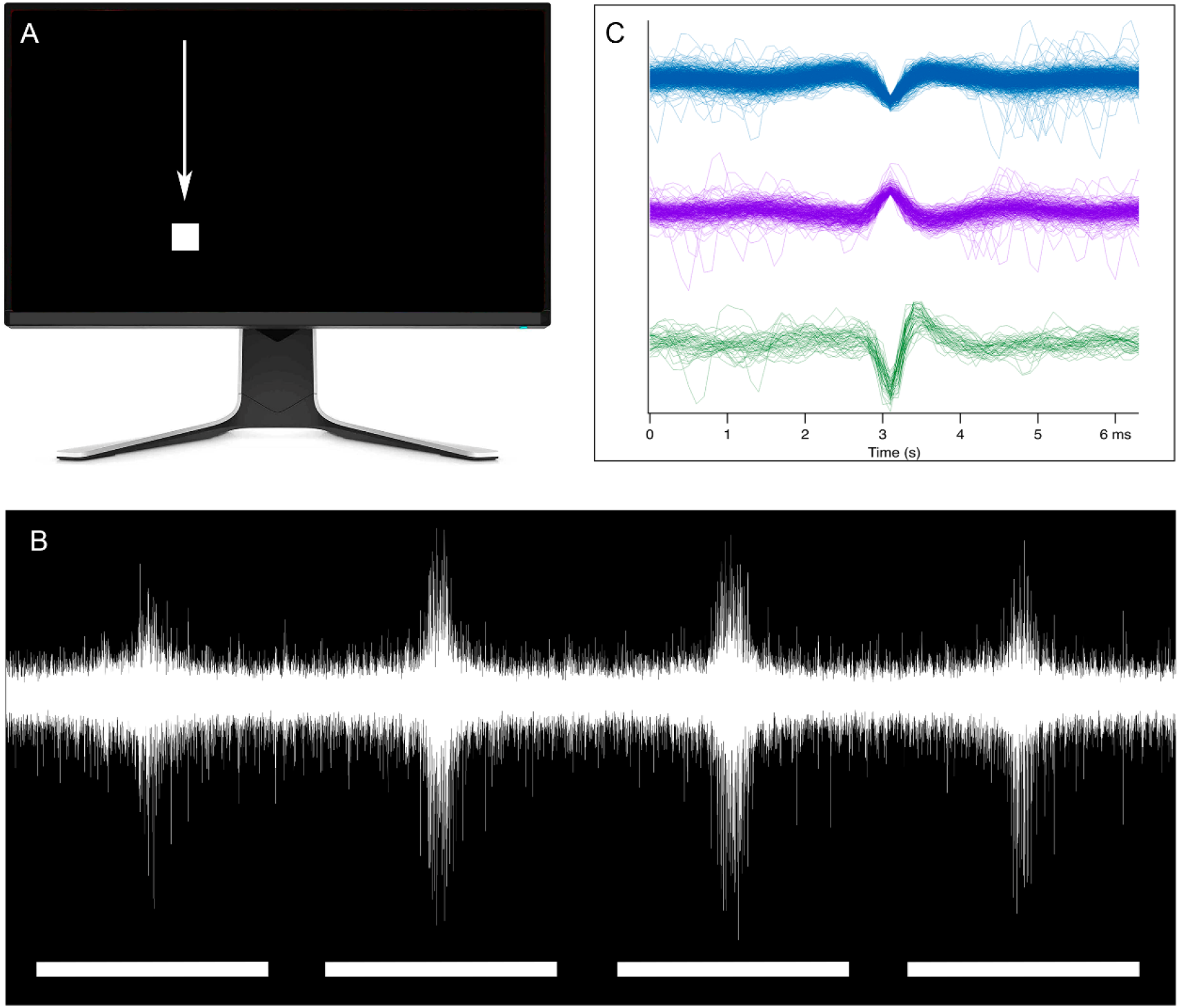
Visual stimulation, extracellular recording, and spike sorting. **(A)** Visual stimuli used to characterize RF size and shape. A small white square (∼3° diameter) on a black background was swept across the screen monitor from top to bottom with each vertical sweep (27°/s, 4 sec), displaced horizontally by 3°, at 1s intervals, to cover the entire screen. **(B)** Top trace, extracellular responses in the SC. Bottom trace, the visual stimulus on/off pattern as described in **A**. **(C)** Example of 3 different single units isolated from recordings.

#### Drifting gratings

Stimuli consisted of full-field, vertical, sinusoidal gratings drifting from temporal to nasal on the LCD monitor. Frequency, contrast and temporal frequency parameters were chosen as in Niell and Stryker (2008). The sinusoidal gratings were presented over 4 s with a 2 s interstimulus interval at a constant temporal frequency (1.5 Hz). All combinations of spatial frequency (0.01, 0.02, 0.04, 0.08, 0.16, 0.32 cycles/deg) and contrast (0.0625, 0.125, 0.25, 0.5, 0.75, 1) were presented in a pseudo-random manner. The stimulation protocol was repeated 5 times for each recording location/depth.

#### Dark looming object

We presented a looming object consisting of a small, black circle (2° diameter) on a gray background, expanding to 20° in diameter over 250 ms, followed by a 250 ms presentation at its maximum size (Yilmaz and Meister 2013).

#### Dark static object

As a control for the looming experiment, we presented a 20° diameter static black circle on a gray background for 500 ms. The looming and static object stimuli were both presented 15 times with 2 s inter-stimulus intervals.

### Data processing

The data were band pass filtered (400 to 5000 Hz) by applying a Finite Impulse Response (FIR) filter with a Hamming window (Choi *et al*. 2006). The background noise was estimated using an adaptive filter algorithm (Biffi *et al*. 2010). Events that crossed a threshold of 3.8 times the background noise were considered as spikes. Spikes were sorted into clusters using the discrete wavelet transform for feature extraction, and coefficients were selected to maximize the clusters’ distance (Letelier and Weber 2000). The clustering of single units was performed using a *k*-means algorithm with optimal *k*-values obtained by the silhouette cluster analysis (Rousseeuw 1987) (see **Fig. 1B-C**).

### Characterization of receptive field properties

#### Receptive field size

The RF area was assessed for each single unit by binning the data into 100 ms intervals, subtracting the spontaneous firing rate, and fitting a two-dimensional (2D) Gaussian curve to the average of the firing rate across trials:

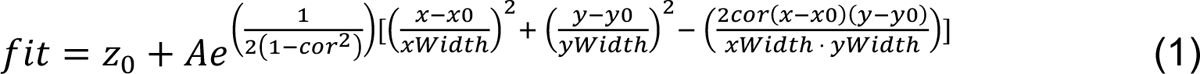

where *z_0_* is the matrix baseline (minimum value of firing rate after the subtraction of the spontaneous firing rate), *A* is the maximal response amplitude, *x_0_* and *y_0_* are the horizontal and vertical positions of the RF’s center, *xWidth* and *yWidth* are the radii of the RF, and *cor* is the correlation coefficient between two correlated Gaussian random variables. The RF areas obtained for each single unit were subsequently categorized into depth ranges, using the ranges defined in Table 1.

**Table 1.** Categorization of depth ranges. Visual responses were systematically recorded from neurons with RFs located within the frontal visual field, at intervals of 100 µm along transects across the SC layers. Neuronal responses were categorized into seven distinct depth ranges according to the depth at which they were recorded.

| Depth range ($\mu\text{m}$)* | SC layer | Depth range |
| --- | --- | --- |
| < 400 | SGS | 1 |
| (400, 500) | SGS | 2 |
| (501, 600) | SO | 3 |
| (601, 750) | SO | 4 |
| (751, 950) | SGI | 5 |
| (951, 1100) | SGI | 6 |
| (1101, 1300) | SGI | 7 |

#### Spatial frequency tuning and contrast responses

Single unit responses were binned in 100 ms intervals and spontaneous firing rate was subtracted. Spontaneous firing rate was obtained from one second interstimulus intervals ranging from 0.5-1.5 s after each stimulus presentation. Units considered in the analysis showed a mean evoked firing rate ≥ 3 Hz after the spontaneous firing rate subtraction. Tuning curves for spatial frequency (SF) were built using mean SF responses at the contrast where the maximum response was evoked, and by fitting the data using a single Gaussian function:

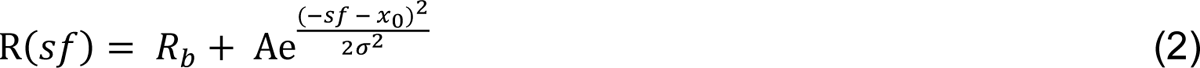

where *R_b_* is the baseline of the curve, A is the response’s amplitude, x_0_ is the position of the center of the peak, and σ is the standard deviation. All variables were optimized by finding the minimum of the unconstrained multivariable function using a derivative-free method from MATLAB-fminsearch (Hendrikje *et al*. 2013; Camillo *et al*. 2020). Neurons with *R^2^* ≤ 0.7 in the Gaussian fitting function were not considered further for analysis. The values for preferred spatial frequency were obtained from each single unit and were classified as low-pass, band-pass or high-pass according to the specific spatial frequency tuning they displayed. *Low-pass neurons* responded to low spatial frequencies (i.e., 0.01 cpd) with a peak firing rate larger than half of the peak’s activity amplitude and showed smaller (less than half of the peak’s amplitude) or non-responses to higher spatial frequencies. *Band-pass neurons* showed a bell shape response with a maximum at its preferred SF while having low firing rates (lower than half of the peak’s response amplitude) for low *sf* (low cut-off) and for high SF (high cut-off). Finally, *high-pass neurons* responded to higher spatial frequencies with firing rates larger than half of the peak response (i.e., > 0.24 cpd). For those neurons showing a band-pass curve we calculated the log_2_ bandwidth (high cut-off frequency/low cut-off frequency) to determine how narrow or wide their tuning was (Heimel *et al*. 2005). We also evaluated the linearity of the responses to the drifting grating stimuli at the preferred spatial frequency by calculating the F1/F0 ratio, with F0 being the mean response and F1 the response modulation at the drifting frequency. Both values were calculated by applying the discrete Fourier transformation (Niell and Stryker 2008; Wang *et al*. 2010). Cells showing a F1/F0 ≥ 1 were classified as linear or simple, whereas those showing a F1/F0 < 1 were classified as non-linear or complex. Single unit responses to different contrasts were obtained by using the spatial frequency value exhibiting the maximum response. The mean responses were fitted using a Naka-Rushton function (Naka and Rushton 1966):

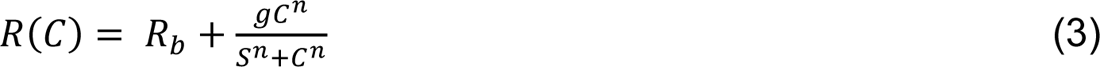

where *R_b_* is the baseline of the curve, *g* the gain, *S* the contrast value at the mid saturation point, *n* the exponent, and *C* the contrast. From the fitted curve we obtained: i) the contrast value at half the peak response (C_50_), which is a measure of contrast response linearity (Heimel *et al*. 2005) and ii) the maximum slope of the fitted curve (maximum rate minus the normalized spontaneous rate), which is a measure for non-linear responses to contrast (Heimel *et al*. 2005).

#### Responses to dark looming objects

Single unit responses to looming and static objects were binned in 10 ms intervals before, during, and after the stimulus presentation. We calculated the neuronal response by first obtaining the spontaneous firing rate for the 300 ms before the stimulus presentation, then we averaged the responses of the 15 repetitions of each kind of stimulus (looming and static objects) and subtracted the spontaneous firing rate. Raster-plots, however, represent the response of a neuron (firing rate minus the spontaneous firing rate) for each trial. To assess the onset time (latency) and peak response time to the object presentation, z-scores were calculated for each bin taking 300 ms before and 500 ms after each stimulus presented (t_stim_ = 0). To ensure the specific response to our visual stimulation we chose a strict threshold; only neurons with a maximum response ≥ 5 z-scores were considered in the analysis. The onset time of the response was the first bin at which the response exceeded 2 z-score values and continued increasing until it reached the peak response. The peak time corresponds to the time of the maximal response between the onset of the stimulus and 250 ms (time for the looming object to reach maximum size).

## RESULTS

### Cytoarchitecture of the degu SC

The mammalian superior colliculus is a laminar structure composed of 7 main layers, namely, stratum zonale (SZ), stratum griseum superficiale (SGS), stratum opticum (SO), stratum griseum intermediale (SGI), stratum album intermedium, (SAI) stratum griseum profundus (SGP) and stratum album profundus (SAP). This latter layer separates the SGC from the underlying periaqueductal grey (PAG). In agreement with previous findings (Vega-Zuniga *et al*. 2013; Deichler *et al*. 2019), we found that SC in degus is a large and highly differentiated structure, in which all layers are well-defined and readily identifiable in Nissl preparations (**Fig. 2A, A’**). As in other mammals (e.g., mouse (Basso and May 2017; Ito and Feldheim 2018), cat (May 2006), tree shrew (Albano *et al*. 1979) and Galago (Balaram *et al*. 2011), but see Major *et al*. (2003)), the SGS in degus appears to contain two subdivisions, upper (uSGS) and deep (dSGS), the former being composed of smaller and less densely packed cells than the latter. Another striking feature is that the SGI in degus appears composed of three subdivisions-upper (uSGI), intermediate (iSGI) and deep (dSGI)-that are readily distinguishable by their differences in cellular density (**Fig. 2A)**, these subdivisions have also been also observed in rats and mice (Wiener 1986; Helms *et al*. 2004).

**Figure 2.**
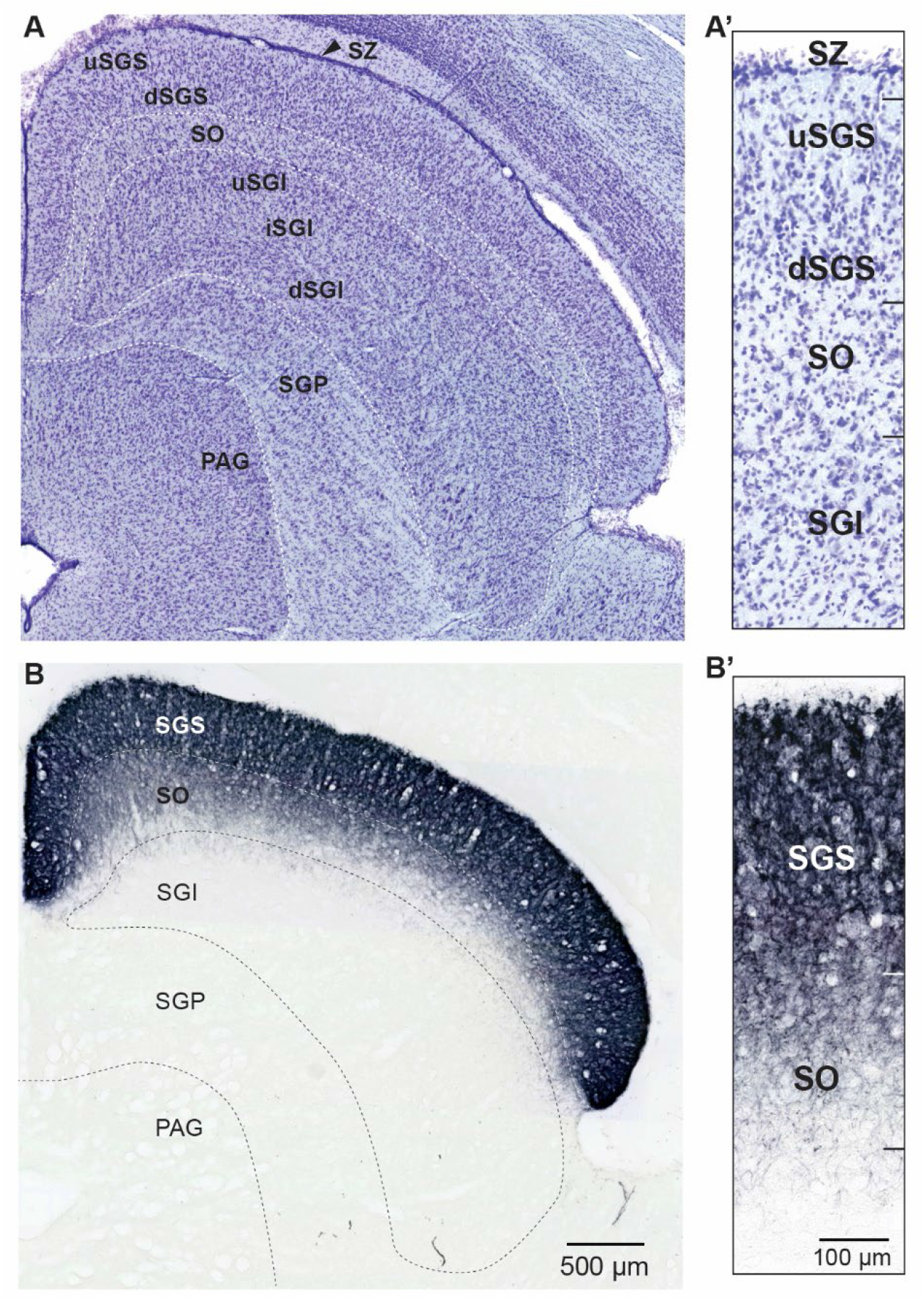
**(A)** Photomicrograph of a Nissl-stained coronal section showing the subdivisions of the degus’ SC: stratum griseum superficiale (SGS) upper (uSGS) and deeper (dSGS), stratum opticum (SO), stratum griseum intermediale (SGI) upper (uSGI), intermediate (iSGI) and deeper (dSGI), the stratum griseum profundus (SGP), and PAG, periacueductal gray. **(A’)** Detail of a Nissl-stained coronal section of SC laminae. **(B)** Anterograde labeling of retinotectal projections after a CTB injection into the contralateral eye. **(B’)** Detail of the retinotectal projection into the different layers of the degus’ SC.

To determine the extent of retinal innervation of SC, we made retinal CTB injections into the vitreous humor. We found that the SC is the largest and most densely innervated retinal target in the degus visual system (**Fig. 2B**). Retinal projections to the degu SC originate almost exclusively from the contralateral eye (Vega-Zuniga *et al*. 2013) and thus we did not quantify the ipsilateral projections. The contralateral retinal projections enter the SC through the SO and terminate very densely in the SZ and in both SGS subdivisions, and more sparsely in the SO. However, the dorsal portion of the SO receives denser terminal arborizations than its ventral part (**Fig. 2B, 2B’**). Retinal projections were not present in the deeper layers of the SC, SGI, and SGP.

### Receptive field properties in the degu SC

To characterize the RF properties in the Chilean degu (*Octodon degus*), we presented different visual stimuli, as detailed in the methods section. Our main objective was to characterize the RF properties within the retino-recipient layers, specifically the SGS and SO, that are part of the superficial SC (sSC). Additionally, we assessed whether neurons in the degu’s SC exhibited the canonical organization of visual RFs characterized by smaller RF sizes in more superficial layers compared to deeper layers, including the SGI.

#### Receptive field size and shape vary with depth in SC

As a first step to characterize the RF properties of SC neurons in degus, and to compare our findings with previous work in hamsters (Carrasco *et al*. 2005) and mice (Fernández-Aburto *et al*., in prep), we recorded from neurons with RFs located in the frontal visual field, and determined how their size and shape varied across the different SC layers (from SGS to SGI). Illustrations of the size and shape of receptive fields for two single units recorded from the SGS and SGI, respectively, are presented in **Figures 3A** and **3B**. Consistent with previous findings, degus showed the canonical organization of visual RFs in the SC (Humphrey 1968), with RF sizes of single units increasing significantly with depth from the SGS to the SGI layer (ANOVA *F*_[6,32]_: 5.58, *p* = 0.00048; **Fig. 3C**). The Tukey HSD (honestly significant difference) post-hoc test revealed that the RF areas of single units recorded from the SGI depth range **7** (see **Table 1**) were significantly larger than RF areas of single units from the SGS depth range **1** (*p* = 0.0006) and significantly larger than RF areas recorded from units in the SGS depth range **2** (*p* = 0.003). RF areas of single units from SGI depth range **7** were also significantly larger than RF areas of the upper SO depth range **3** (*p* = 0.003; **Fig.3C**). In addition, we compared RF sizes between the SC layers combined (SGS, SO and SGI), and found a significant difference (ANOVA *F*_[2,36]_: 12.46, *p* < 0.0001; **Fig. 3D).** RF areas from SGI neurons (area size range: 42.84-162.8 deg^2^) were significantly larger (Tukey HSD, *p* = 0.0001) than RF areas from SGS neurons (area range: 19.8 – 90.97 deg^2^), and significantly larger (Tukey HSD, *p* = 0.0018) than RF areas from SO neurons (area range: 30.35 – 86.01 deg^2^). RF sizes from SGS neurons and SO neurons were similar (Tukey HSD test, *p* = 0.36).

**Figure 3.**
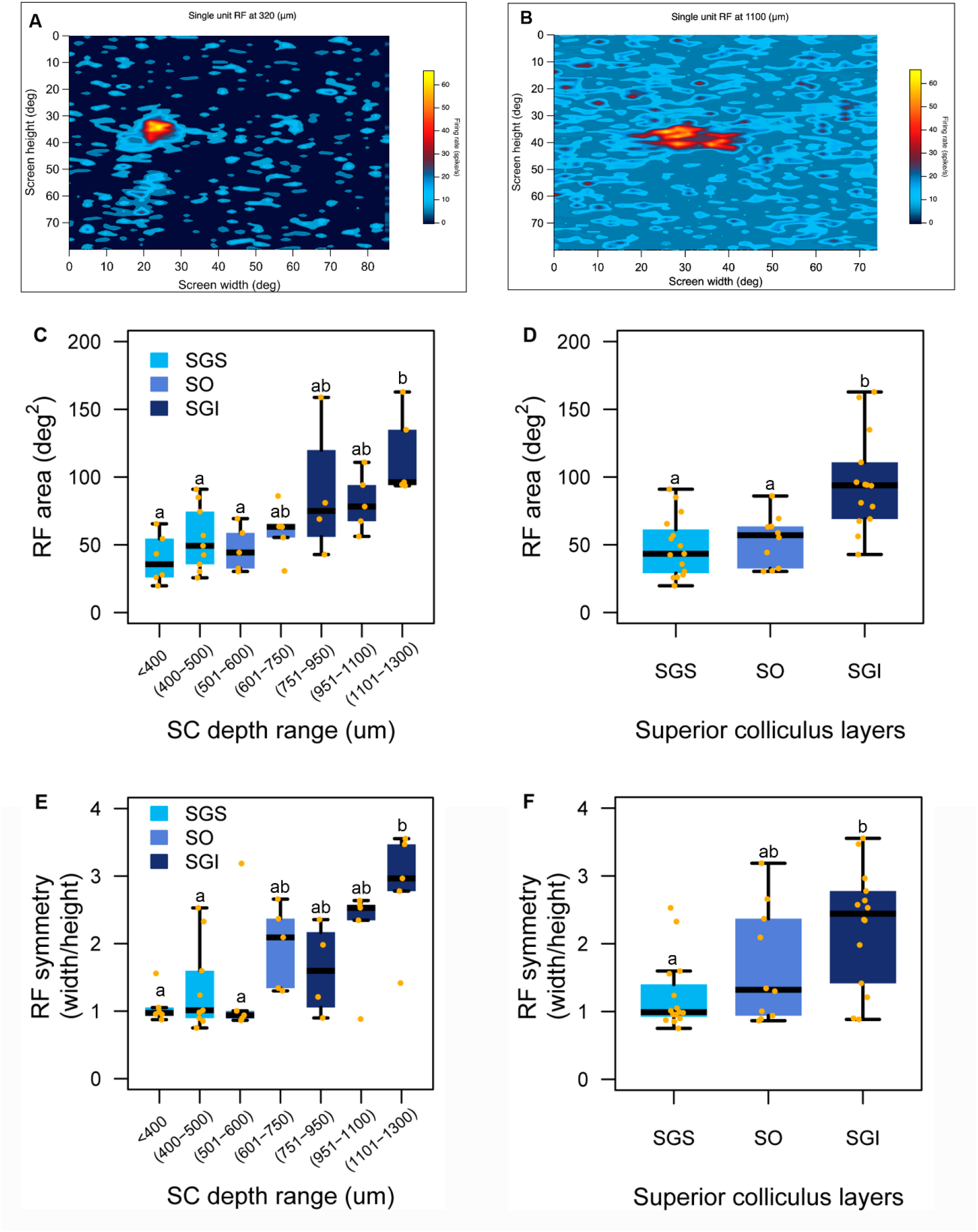
Receptive field size and shape change across SC layers in degus. **(A)** Heat map showing the RF of a single unit isolated at 320 µm depth in the SGS mapped by visual stimulation as in Fig. 1A. **(B)** Heat map showing the RF of a unit at 1100 (µm) in the SGI mapped with the same stimulation, at the same recording site (stereotaxic coordinates) as in **A**, but deeper**. (C)** Boxplot of RF areas (in deg^2^) across depths ranges from the SGS to the SGI. Boxplots show standard interquartile range (IQR) and median value (black central line), yellow dots are the individual data points. RFs increase in size with depth (ANOVA: *p* < 0.01). **(D)** Boxplot of RF areas across SC layers (SGS, SO and SGI). **(E)** Boxplot of RF shape expressed as the width/height ratio at different SC depth ranges. RFs widen along the horizontal axis with depth (ANOVA: *p* <0.01). **(F)** Boxplot of RF shape at different SC layers. Different letters above each group in B, C, D, and F represent statistically significant differences (Tukey post-hoc test, see text for exact *p*-values).

Studies on rat and monkey SC have shown that there is a change in RF shape across SC layers, going from near circular RFs in the superficial layers to RFs elongated along the horizontal axis in the deeper layers (Humphrey 1968). To investigate whether degus also exhibit an alteration in receptive RF shape (i.e., circular vs. oval shape), we compared the ratio between the horizontal and vertical axes of the RFs across the SC depth ranges. A ratio close to 1 indicates a symmetrical RF with a circular shape, whereas deviations from this value indicate an elongation of the horizontal axis. Similar to the findings reported by Humphrey (1968), RF symmetry changes significantly between depth ranges; RFs from deeper layers are more elongated along the horizontal axis than RFs from upper layers (ANOVA *F*_[6,32]_: 4.04, *p* = 0.004; **Fig. 3E**). The RFs of neurons from SGI depth range **7** were more elongated along the horizontal axis (i.e., asymmetrical) than the circular RF shape of neurons from SGS depth range **1** (*p* = 0.004) and were also more elongated than the RF shape of neurons recorded from the SGS depth range **2** (*p* = 0.01). Likewise, neurons from SO depth range **3** showed RF shapes that are significantly more circular than the elongated RF shapes of neurons from SGI depth range **7** (*p* = 0.037; **Fig. 3E**). Comparisons focusing on the differences between SC layers as a whole rather than the depth level confirmed that as recordings sampled the lower layers, RFs shapes became more elongated along the horizontal axis (ANOVA *F*_[2,36]_: 6.69, *p* = 0.003; **Fig. 3F**). The RFs of SGS neurons were significantly more symmetric (i.e., circular) than RFs of SGI neurons (Tukey HSD, *p* = 0.002). The shape of the RFs from SO neurons was statistically the same as the RF shape of single units from the SGS and SGI, suggesting a tendency of SO neurons to have elongated RFs.

#### Spatial frequency tuning

Sinusoidal gratings of different spatial frequencies (SF) and contrasts (**Fig. 4A**) have been widely used to characterize RF properties of different visual brain structures in several animal species (Jassik-Gerschenfeld and Hardy 1979; Issa *et al*. 2000; Heimel *et al*. 2005). In the case of degus, we found that 47% (228/485) of SGS neurons that were recorded responded to sinusoidal grating stimuli. Of those, 58.3% (133/228) were tuned to specific combinations of SF and contrast (see below). According to their spatial frequency tuning, we classified neurons as low-pass, band-pass or high-pass (see Methods). Most SC neurons were band-pass (60.9%), or low pass (36.8 %); only a few were classified as high-pass neurons (2.3%; see Figures **4B** through **4E’**). Even though the median value of SF preference across all three cell types (low-pass, band-pass and high-pass) was 0.04 cycles per degree (cpd), we found a subset of band-pass neurons (18%) tuned to a higher SF (0.24 cpd) (**Fig. 4F**).

**Figure 4.**
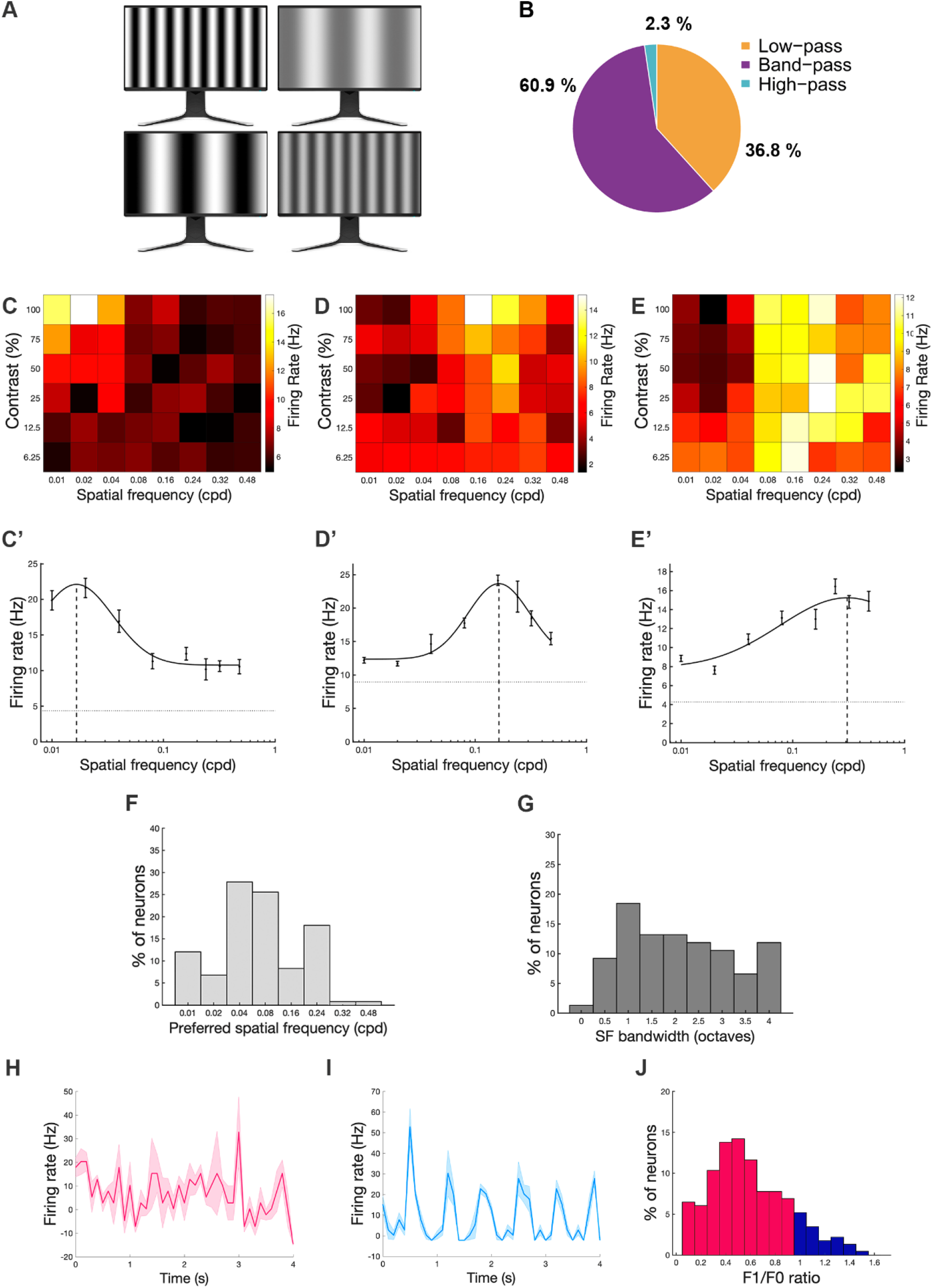
Single units show selectivity to spatial frequency. sSC neurons have a narrow bandwidth, and a small proportion of them (14%) respond linearly to sinusoidal gratings. **(A)** Illustration of the drifting grating stimulus types. Spatial frequency and contrast were varied in a pseudorandom manner. **(B)** Percentage of neurons with low-pass, band-pass or high-pass selectivity to different spatial frequencies (n = 128). **(C-E)** Response matrix (instantaneous firing rate, in Hz) of three different neurons to all contrast and spatial frequency combinations. **(C’-E’)** Examples of low-pass **(C’)**, band-pass **(D’)**, and high pass **(E’)** tuning of the same single neurons as in **C-E**. **(F)** Bar plot of spatial frequency preference for the population. Most units are tuned to 0.04 or 0.08 (cpd), and a smaller group is tuned to higher spatial frequency (0.24 cpd). **(G)** Bar plot of bandwidth showing that most neurons (>50%) are tuned to a certain spatial frequency with narrow bandwidths. **(H-I)** Solid lines are the mean firing rate (100 ms bins) and shaded regions represent mean ± sd. **(H)** Example of a single unit responding non-linearly to 4 seconds of stimulation with a sinusoidal grating (linearity index F1/F0 < 1, where *F1* is the response at the drift frequency, and *F0* is the average firing rate). **(I)** Example of a different single unit responding in a linear manner to sinusoidal gratings (F1/F0 >1). **(J)** Histogram of the linearity index across the entire population of single units (n = 232) showing neurons responding linearly to sinusoidal gratings (red bars) and neurons with non-linear responses to sinusoidal gratings (blue bars).

To further assess the specificity of SF tuning in band-pass neurons, we quantified the tuning curves’ bandwidths (the range of SF to which a band-pass neuron responds). Interestingly, 41% of band-pass neurons were tuned to their preferred SF, with bandwidths ranging between 0.5 and 2.0 octaves (**Fig. 4G**). Additionally, we assessed the extent to which spatial frequency modulates the activity of degus’ SC units by calculating the linearity index *F1/F0* (where *F1* is the modulation of the response (firing rate) at the drift frequency of the stimulus, and *F0* is the average firing rate of the response, see Methods) (De Valois *et al*. 1982). We found that most SGS neurons (86%) responded non-linearly to their preferred SF (*F1/F0* < 1) **(Fig. 4H)**, and the rest (14%) responded linearly (*F1/F0 > 1*) (**Fig. 4I)**. The distributions of linear/non-linear responses for the neuronal population is shown in Figure **4J**.

#### Contrast tuning

Overall, SGS neurons increased their response levels with increased contrast. However, there were neurons that showed a linear response to contrast without saturating their response (53.3%; **Fig. 5A**), and others (46.7%) that exhibited different degrees of saturation at higher contrast (**Fig. 5B-C**). Interestingly, such saturating responses are characterized by high contrast gain at low contrasts and low contrast gain at high contrasts (Alitto *et al*. 2019). We further characterized linear and saturating responses by quantifying the contrast at half of the maximum response (C_50_ value) and the relative maximum gain (RMG), i.e., the maximum slope of the normalized contrast response curves (Heimel *et al*. 2005). At the population level, 50% of the neurons showed that the contrast at half of the maximum response (C_50_ value) was 48%, indicating a linear response to contrast (**Fig. 5D**). On the other hand, neurons that show saturating responses can have high (C_50_ < 30, 36% of neurons) or moderate (C_50_ between 30 and 40, 17% of neurons) saturation of their responses to contrast. Additionally, the median value of RMG was 1.47 (**Fig. 5E**), indicating slight non-linearity, compared to non-linear responses to contrast in the squirrel’s V1 with a median RMG value of 2.8 (Heimel *et al*. 2005). Only 22/135 neurons (16.3%) had a RMG of approximately 1 (between 0.8 and 1.4) that we considered as a linear behavior, whereas 28/135 showed RMG values ≥ 3 (20.7%) and were thus considered non-linear. Interestingly, 50% of neurons (170/477) showed RMG values between 1.4 and 1.5, which we interpreted as weak non-linear responses. Although both measures, C_50_ and RMG, are indicators of linearity, the RMG is less dependent on the fitting process (Heimel *et al*. 2005). Still, both measures suggest that the degus SGS has neuron types that respond in a linear and a nonlinear fashion.

**Figure 5.**
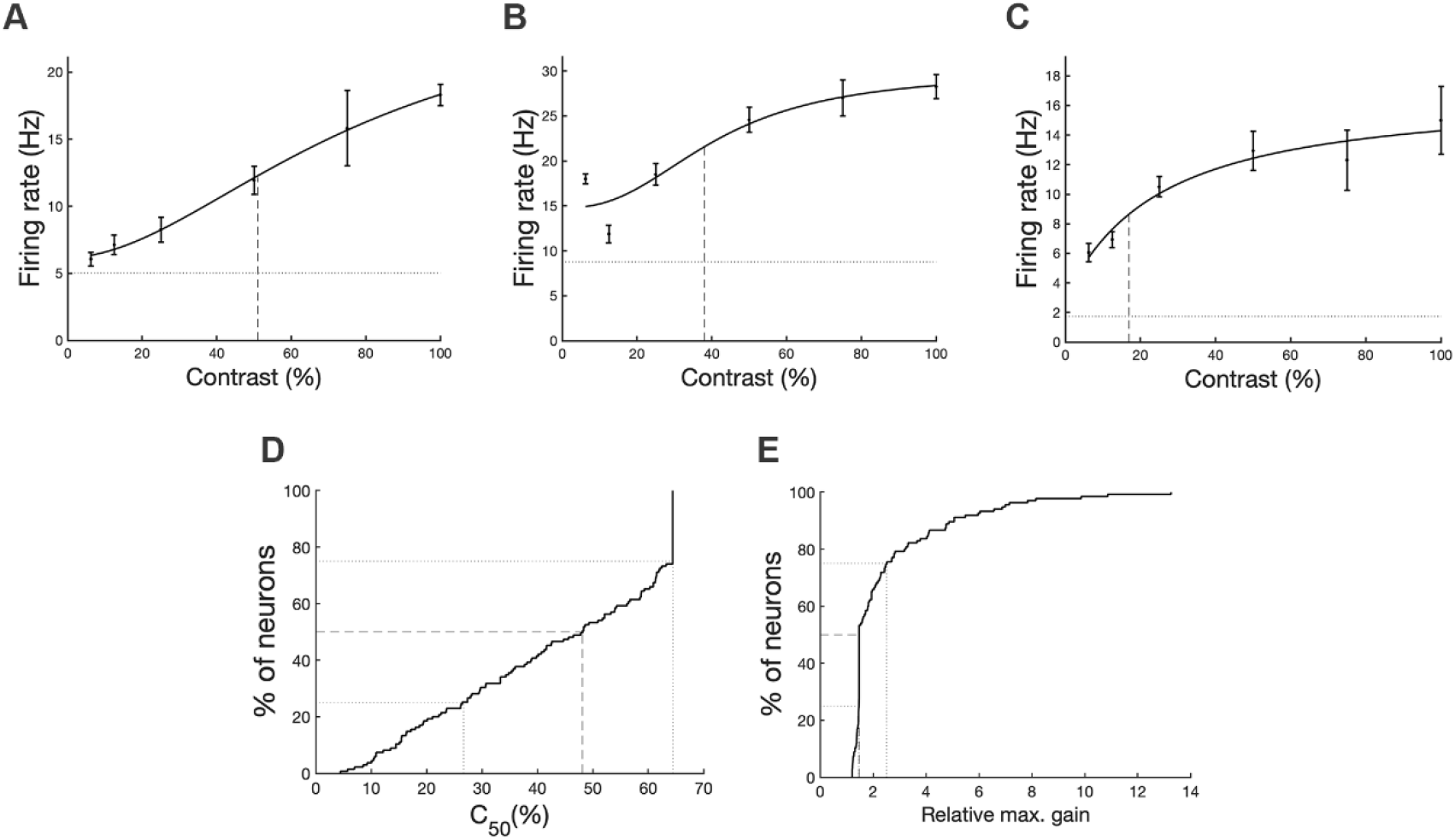
SC Single units show linear or non-linear responses to contrast. Example of three different single units’ tuning to contrast, ranging from linear **(A)** to non-linear **(B and C).** All three curves are fitted with a Naka-Rushton function (black line). The dotted line is the spontaneous firing rate (Hz) and the dashed line is the C_50_ (the contrast at half the peak response). Note that **(B)** and **(C)** show saturating responses and differ in their C_50_ values (∼40 and ∼20, respectively). **(D)** Cumulative histogram of C_50_ distribution (N=135), with the median C_50_ = 48. **(E)** Cumulative histogram (N=135) of the relative maximum gain (see Methods).

#### SGS neurons respond in a continuously increasing way to looming

To evaluate the units’ responses to looming stimuli, we presented 15 repetitions of a dark disc that increased in size from 2 to 20 deg in 250 ms and stayed ON for another 250 ms. As a control we presented a static dark disc (20°) for 500 ms to address the specificity of the response to looming (**Fig. 6A and A’**). Neurons exposed to the looming object continuously increased their firing rate as the object increased in size, followed by a decrease of the firing rate when the object no longer increased in size. A small OFF response can be observed (**Fig. 6B and 6C)**. Neurons exposed to the static object, on the other hand, showed a typical ON-OFF response (**Fig. 6B’ and 6C’**). Of a total of 251 responsive neurons, 146 responded to both stimuli, 58 only to looming, and 47 only to the static object. Interestingly, the onset time of the response to both stimuli did not differ significantly (paired *t*-test, *p* = 0.3), showing that neurons start their response at the same time irrespective of the visual object shown (**Fig. 7A**). The difference between the response to the looming and to the static object can be observed in other parameters-in peak time (**Fig. 7B**) and maximal firing rate after the initial response (**Fig. 7D**). With the looming parameters used, neurons exposed to the looming take longer to reach their maximal response (paired *t*-test, *p* = 0.0066) and show significantly higher firing rates (paired *t*-test, *p* < 0.0016) than when exposed to the static object.

**Figure 6.**
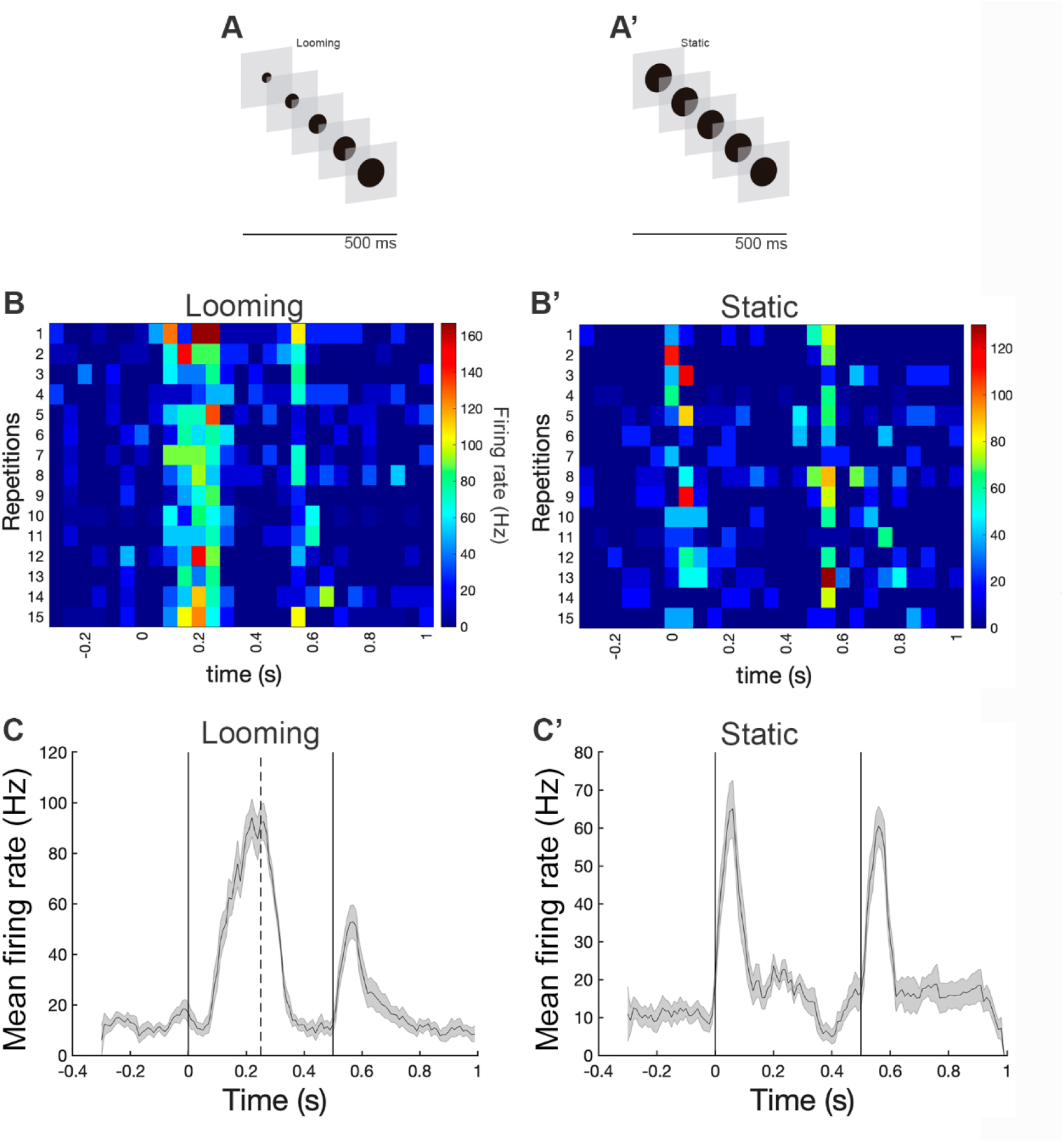
Single units show a continuous buildup of response level to looming stimuli but an ON-OFF response to static stimuli. **(A)** Diagram of visual stimulation. The looming object increased in size from 2° to 20° over 250 ms, followed by 250 ms at the 20°size. **(A’)** The static object was 20° in diameter and was presented for 500 ms. **(B-B’)** Heatmap showing instantaneous firing rates of a single unit responding to 15 consecutive presentations of a looming object **(B)** or of a static object **(B’). (C-C’)** PSTH of the same single unit showing the mean firing rate for responses to the looming **(C)** and static objects **(C’)** across the 15 trials (mean ± SE).

**Figure 7.**
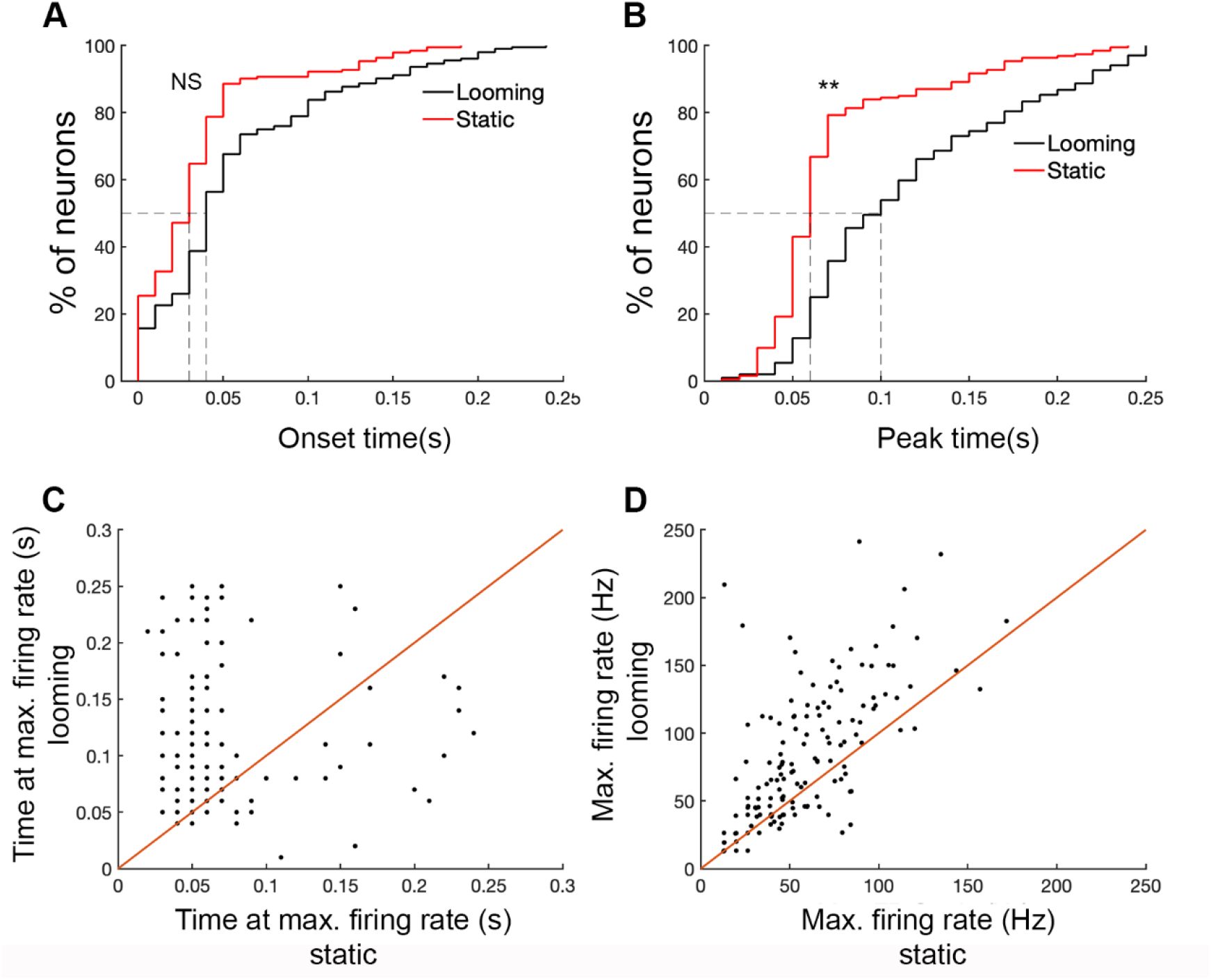
Neurons respond differentially to looming and static objects. **(A)** Cumulative probability for onset time comparing both visual stimulus types. Single units responded with similar onset time to both types (static object (red), looming object (black)). **(B)** Cumulative probability for the timing of peak activity comparing responses to looming (black) and static visual stimuli (red). Single units reach their peak activity significantly faster with the static object than with the looming object (p ≥ 0.05). **(C)** Time at maximal firing rate of neurons responding to both the looming and to the static object; note that most neurons fall above the identity line (slope = 1.0; red) indicating that their maximal firing occurs later in time when exposed to looming compared to the static object. **(D)** Maximum firing rate to each visual stimulus of neurons that responded to both stimuli. Interestingly, neurons show higher firing rates when exposed to looming stimuli.

## DISCUSSION

It is well known that mature visual properties as well as visual system development differ between mammalian species in a way that is related to the role played by vision in their behavior. Interestingly, most of the recent publications about visual system development and function come from a single model species, the lab mouse *(Mus musculus)*. Lab mice are altricial, nocturnal, and feature a poorly developed visual system when compared with diurnal rodents such as squirrels, guinea pigs and some other New World rodents. Thus, mouse-centered studies may have biased the current understanding of rodent visual system function and its development. As a starting point to counterbalance this bias, we characterized the visual properties of SC neurons in the adult *Octodon degus* in the present study. Degus, contrary to mice and other typical lab rodent species such as rats and hamsters, are diurnal (Fulk 1976) and precocial (Reynolds and Wright 1979; Long and Ebensperger 2010). Furthermore, degus represent a phylogenetic group (suborder Hystricomorpha) apart from old world rodents such as rats and mice (suborder Myomorpha) or squirrels (suborder Sciuromorpha). As we predicted, we found that *O. degus’* SC is a well-developed visual structure, featuring visual properties comparable to that of highly visual rodents such as squirrels (Van Hooser *et al*. 2003) and gerbils (Baker and Emerson 1983). We conclude that, given its precocial and diurnal habit, *O. degus* is especially well-suited as an animal model for studies of visual system development.

### Cytoarchitecture of the superior colliculus in *Octodon degus*

Our anatomical results readily confirmed that degus’ SC features a laminar differentiation comparable to that of other diurnal mammals, such as the squirrel (Major *et al*. 2003; Fredes *et al*. 2012). In particular, the extensive laminar differentiation of the SGI suggests that in degus the SC serves a stronger and more complex visuomotor role than in other rodents, a conjecture worthy of further studies. In addition, the results of our retinal injections confirm previous studies (Vega-Zuniga *et al*. 2013) reporting that degu SC receives massive and highly ordered retinal projections. As in the previous studies, we found that retinal afferents enter the contralateral SC through the SO to form dense terminal domains in the stratum griseum superficiale (SGS) and stratum zonale (SZ), and sparser arborizations in the stratum opticum (SO). As expected from studies of other species, retinal projections were not present in the deeper layers of the SC (dSC), namely the stratum griseum intermediale (SGI) and the stratum griseum profundus (SGP).

### Receptive field size

We found that the degu’s SC has a laminar segregation of RF sizes, with smaller RFs located in superficial layers, and larger RFs in deeper layers. This seems to be a canonical property of the SC - *aka* optic tectum-across vertebrate taxa, in that it has been reported not only in other rodents, such as mice, rats and hamsters (Humphrey 1968; Rhoades and Chalupa 1976; Ito *et al*. 2017), but also in Carnivore species (cats and ferrets (Brecht *et al*. 1999; Stitt *et al*. 2013), primates (Humphrey 1968; Cynader and Berman 1972) and birds (Knudsen 1982; Marín *et al*. 2005)). This change in RF size across layers correlates with the different morphological characteristics of neurons located in each layer. In mammals, narrow field and stellate neurons located in the upper SGS (uSGS) have narrow dendritic trees and small RFs (Mooney *et al*. 1985; May 2006; Hoy *et al*. 2019), whereas wide field and horizontal cells, located in the deeper SGS (dSGS), have large dendritic trees (∼500-800 μm) along with very large RFs (∼300-1000 deg^2^) (Major *et al*. 2000; Fredes *et al*. 2012; Gale and Murphy 2014). In the dSC (SGI and SGP), neurons that have large dendritic trees also exhibit large RFs, most of them being premotor neurons that receive convergent inputs from neurons located in the superficial layers (Isa and Hall 2009).

Superior colliculus RF sizes are highly variable across rodent species. In hamsters the SGS RFs are quite large (ranging from 169 deg^2^ to ∼295 deg^2^ [19.4 ± 0.56° diameter] (Carrasco *et al*. 2005; Balmer and Pallas 2015). In mice, on the other hand, contrasting results have been reported. Some studies report large RF sizes (average 165 ± 21 deg^2^ (Chandrasekaran *et al*. 2005; Ito *et al*. 2017) but others report smaller values (107.5 ± 14.9 deg^2^, (Wang *et al*. 2010)). These differences may depend on the visual stimulation method used. Ito *et al*. (2017) used a 10° flashing-spot but Wang *et al*. (2010) used a 5° flashing square. In the present study we used a small (3°), moving bright square as a visual stimulus, which is bigger than that used in previous studies of hamsters (1°) (Carrasco *et al*. 2005; Mudd *et al*. 2019). Nevertheless, despite methodological differences, degus seem to have the smallest RF sizes reported in rodents for the sSC so far, with area sizes that never surpassed 100 deg^2^ (ranging from 19.8 deg^2^ to 74.4 deg^2^). It is noteworthy that degus are diurnal animals and possess a cone-rich retina (Chávez *et al*. 2003; Jacobs *et al*. 2003; Vega-Zuniga *et al*. 2013). These features suggest that degus would have a visual acuity higher than that of nocturnal species. The comparatively small size of degu SC receptive fields is indeed consistent with a greater visual acuity, reflecting the diurnal specialization of the visual system in these rodents.

### Response to grating stimuli

In cats, hooded rats, and squirrels, it has been shown that the presence of visual neurons tuned to high spatial frequencies correlates with higher visual acuity (Dean 1981; Issa *et al*. 2000; Heimel *et al*. 2005). In the case of degus, we found that most sSC neurons (> 50%) that responded to gratings exhibited preferences for low/medium spatial frequencies. Interestingly, we found another set of neurons (15%) that were tuned to higher SFs (with a peak of 0.24 cpd). Overall, the SF tuning reported for mouse SC [peak=0.08 cpd, (Wang *et al*. 2010); median= 0.052 cpd, (De Franceschi and Solomon 2018) is lower than what we found in degus, suggesting that, as expected from its diurnal visual specialization, degus have a higher visual acuity than mice. Interestingly, it has been reported that awake mice show an approximately 2-fold higher SF tuning than anesthetized mice (De Franceschi and Solomon 2018)]. Here we only tested deeply anesthetized animals, thus awake degus might show higher SF tuning than we report here.

#### Spatial frequency modulation

In degu sSC, most (86%) of the neurons recorded exhibited non-linear (F1/F0 <1) responses to grating stimuli, thus showing low modulation by the SF presented. However, some neurons (14%) were modulated by the SF (F1/F0 >1). A similar distribution of linear and non-linear neurons was found in mice (Wang *et al*. 2010). Interestingly, neurons showing low or no SF modulation were tuned to higher frequencies, a phenomenon also reported in mice (De Franceschi and Solomon 2018).

#### Contrast sensitivity

We found that almost half (44%) of the neurons responded in a linear, monotonically increasing fashion to increasing stimulus contrast (C_50_ > 45), whereas others showed highly or moderately saturating responses to contrast. It has been proposed that saturated responses to contrast, rather than being the result of biophysical limitations of neurons, are due to an active mechanism controlling the gain of contrast responses (Peirce 2007). Thus, saturating responses to contrast relate to higher sensitivity to subtle contrast differences at low contrast values, which is an important adaptation for active nocturnal vision. In mice most SC neurons exhibit non-linear, saturating responses to contrasts (De Franceschi and Solomon 2018), as expected for nocturnal species. Unlike what has been observed in mice, we found no clear bias toward saturating responses to contrast in the degus’ SC, likely reflecting the diurnal adaptations of their visual system.

### Looming responses

In degus, we found many sSC neurons that showed a progressive increase in their responses to an expanding dark disc that simulated a dark looming stimulus approaching against a lighter background. This is similar to what has been reported in mice and hamsters, in which neurons located in sSC respond reliably to repeated presentation of a dark looming stimulus in their RFs (Mudd *et al*. 2019; Lee *et al*. 2020). Most of the neurons recorded in this study (> 55 %) showed a clear difference in their response properties between looming and static stimuli, which reflects the importance of animals responding differentially to different kinds of stimuli, i.e., a fast-approaching object (e.g., a predator) vs. a large, stationary object. In this regard, we observed that neurons responding to both stimulus types (looming and static objects) had significantly higher firing rates when exposed to the looming object than when exposed to the static control, suggesting that the SC possesses a neural mechanism that allows it to identify a fast-approaching object. This mechanism might be involved in the execution of behaviors in this context. In rodents, looming stimuli have been reported to unleash behaviors modulated by SC such as escaping and freezing (Yilmaz and Meister 2013; De Franceschi and Solomon 2018). In degus, looming stimuli projected from the upper, binocular portion of the visual field elicit escape behavior, which is thought to be triggered by the SC/parabigeminal nucleus interaction (Deichler *et al*. 2020).

## Conclusion

Degus show remarkably specialized visual properties in the superior colliculus compared to commonly studied lab rodents. Our results suggest that degus have diurnal visual characteristics, and a higher visual acuity than other lab rodents. Moreover, degus are precocial, diurnal and belong to a separate clade within the Rodentia order, allowing studies to address from a comparative perspective which visual features are conserved among rodents or other mammals, and which might be determined by their ecological niche. Altogether, we strongly argue that degus are a promising model to study the visual system in general and its development in particular.

## Acknowledgements

We would like to thank Solano Henríquez for his valuable assistance.

## Notes

### Competing Interest Statement

The authors have declared no competing interest.

### Summary of Updates

Improvements in writing and organization. Reorganization of abstract.

